# CellContrast: Reconstructing Spatial Relationships in Single-Cell RNA Sequencing Data via Deep Contrastive Learning

**DOI:** 10.1101/2023.10.12.562026

**Authors:** Shumin Li, Jiajun Ma, Tianyi Zhao, Yuran Jia, Bo Liu, Ruibang Luo, Yuanhua Huang

## Abstract

A vast amount of single-cell RNA-seq (SC) data has been accumulated via various studies and consortiums, but the lack of spatial information limits its analysis of complex biological activities. To bridge this gap, we introduce cellContrast, a computational method for reconstructing spatial relationships among SC cells from spatial transcriptomics (ST) reference. By adopting a contrastive learning framework and training with ST data, cellContrast projects gene expressions into a hidden space where proximate cells share similar representation values. We performed extensive benchmarking on diverse platforms, including SeqFISH, Stereo-Seq, 10X Visium, and MERSCOPE, on mouse embryo and human breast cells. The results reveal that cellContrast substantially outperforms other related methods, facilitating accurate spatial reconstruction of SC. We further demonstrate cellContrast’s utility by applying it to cell-type co-localization and cell-cell communication analysis with real-world SC samples, proving the recovered cell locations empower novel discoveries and mitigate potential false positives.

## Introduction

Spatial-resolved transcriptomic profiling has emerged as a crucial tool for investigating both cellular gene expression and its corresponding positional context. This approach is vital for unraveling complex tissue organizations and intercellular communication that underlies cooperative biological functions. However, the current landscape of spatial transcriptomic (ST) technologies, spanning imaging- and sequencing-based methods ^1^, face several challenges that prevent comprehensive problem resolution. Imaging-based methods, such as SeqFISH+ ^2^, STARmap ^3^, and MERFISH ^4^, have achieved single-cell resolution; however, their gene profiling capabilities are limited to a few hundred predefined genes. Conversely, sequencing-based methods like 10X Visium, Slide-seq ^5^, and Stereo-Seq ^6^ provide the complete transcriptome, albeit at spatial resolutions larger than a single cell. In recent years, single-cell RNA sequencing (SC) has become a widely used technology, and large data resources have been built based on it. Although SC data lacks spatial information, it yields more comprehensive gene profiles than imaging-based ST methods, while also affording higher resolutions than sequencing-based ST approaches. Therefore, researchers have started to explore combining ST and SC to overcome their individual limitations. Various methods have been developed to enhance ST data with SC as a reference, including spot deconvolution ^7, 8^ and gene imputation ^9, 10^. While these methods address the limitations of ST to some extent, the potential of existing vast SC resources remains untapped.

An alternative paradigm has arisen from an inverse perspective, whose aim is to reconstruct the spatial locations of SC data using ST as a spatial reference. Several methods have been proposed, including NovoSpaRc ^11^, which assigns SC cells to predefined locations based on the assumption that physically proximate cells share similar gene expression profiles. Its location assignment is performed based on optimal transport. Although NovoSpaRc does not necessarily require an ST reference, the locations and marker genes derived from ST can be utilized to improve its performance. Subsequently, Tangram ^12^ and CytoSpace ^13^ were introduced to establish mapping relationships from SC to ST. Although both methods minimize the correlation-based cost function inferred from gene expressions, their assumptions and outcomes are different. Tangram considers cell density and gene expression, generating mapping probabilities from SC to ST. CytoSpace generates distinct SC-to-ST predictions, based on inferred cell counts per ST spot and corresponding gene expression. However, Tangram can generate low probabilities for matched SC-ST pairs, and CytoSpace might filter out some SC cells to ensure optimal subsets of samples, potentially leading to incomplete reconstructions. Recently, supervised methods have emerged, such as SageNet ^14^ and CeLEry ^15^. SageNet employs a graph neural network to model the gene interaction network from the ST data and produces an estimated cell-cell distance matrix for the query cells. It has demonstrated accurate and robust performance across distinct samples. However, the preprocessing step of making spatial partitions is dataset-specific and intricate for non-experts. The memory consumption of SageNet also explodes when the number of genes increases. CeLEry represents another supervised deep neural network approach for recovering spatial information of SC data. By ingesting the gene expression inputs, CeLEry was trained by minimizing the loss between the predicted domain layer or spatial coordinates and the truth. It has shown reliable performance in global spatial reconstruction across various species and technologies. However, it does not fully consider the performance of local cell environment reconstruction, which is vital for many downstream tasks.

In this study, we introduced cellContrast, a deep-learning method that employs a contrastive learning framework for spatial relationship reconstruction of SC data. Our fundamental assumption is that gene expression profiles can be projected into a latent space, where physically proximate cells demonstrate higher similarities. To achieve this, we employed a contrastive framework of an encoder-projector ^16^. We built training batches containing positive and negative contrastive pairs based on their truth spatial relationship. We optimized the model using infoNCE loss ^17^, which pulls positive pairs closer and pushes negative pairs apart. During inference, we discarded the projector and used the output of the encoder for spatial relationship reconstruction, based on the principle that higher cosine similarity indicates shorter spatial distance. We benchmarked the performance in multiple settings, including matched samples, independent samples, and independent platforms between training and testing with mouse embryos, human breast, and mouse brain data. Our experiments found that cellContrast outperformed related methods by a large margin and that specifically, local heterogeneity can be well captured. We also demonstrated the power of spatial reconstructed SC data in two downstream tasks: cell-type spatial co-localization and cell-cell communication (CCC). The results showed that with the help of cellContrast, cell-type co-localization patterns were well detected and potential false positive CCC signals were reduced.

## Results

### Overview of cellContrast

The aim of cellContrast is to train an encoder that extracts spatially preserved representations based on the gene expression profiles of ST datasets. The fundamental assumption is that gene expressions can be projected into a feature space where spatially proximate cells exhibit similarity. This trained encoder is then applied to SC samples, enabling estimation of distances between SC samples and between SC and ST samples. We adapted contrastive learning for this purpose (Methods). As illustrated in Figure 1A, our method employs a deep neural network comprising two main components: an encoder for spatial semantic representation extraction and a projection head for loss calculation. Both components were implemented as multi-layer perceptron (MLP). Constructing the training batches containing contrastive pairs is crucial. Given batch size *N*, we first randomly selected *N*/2 cells. For each sample X, we selected one of its *k* nearest neighbors as the positive pair and the rest of *N*-2 cells (the 2 being the X and its random selected positive pair) served as the negative pairs. The projection head generated vectors that were used for infoNCE loss calculation, which encouraged positive pair similarity and negative pair dissimilarity. Model optimization was achieved via SGD.

**Figure 1.**
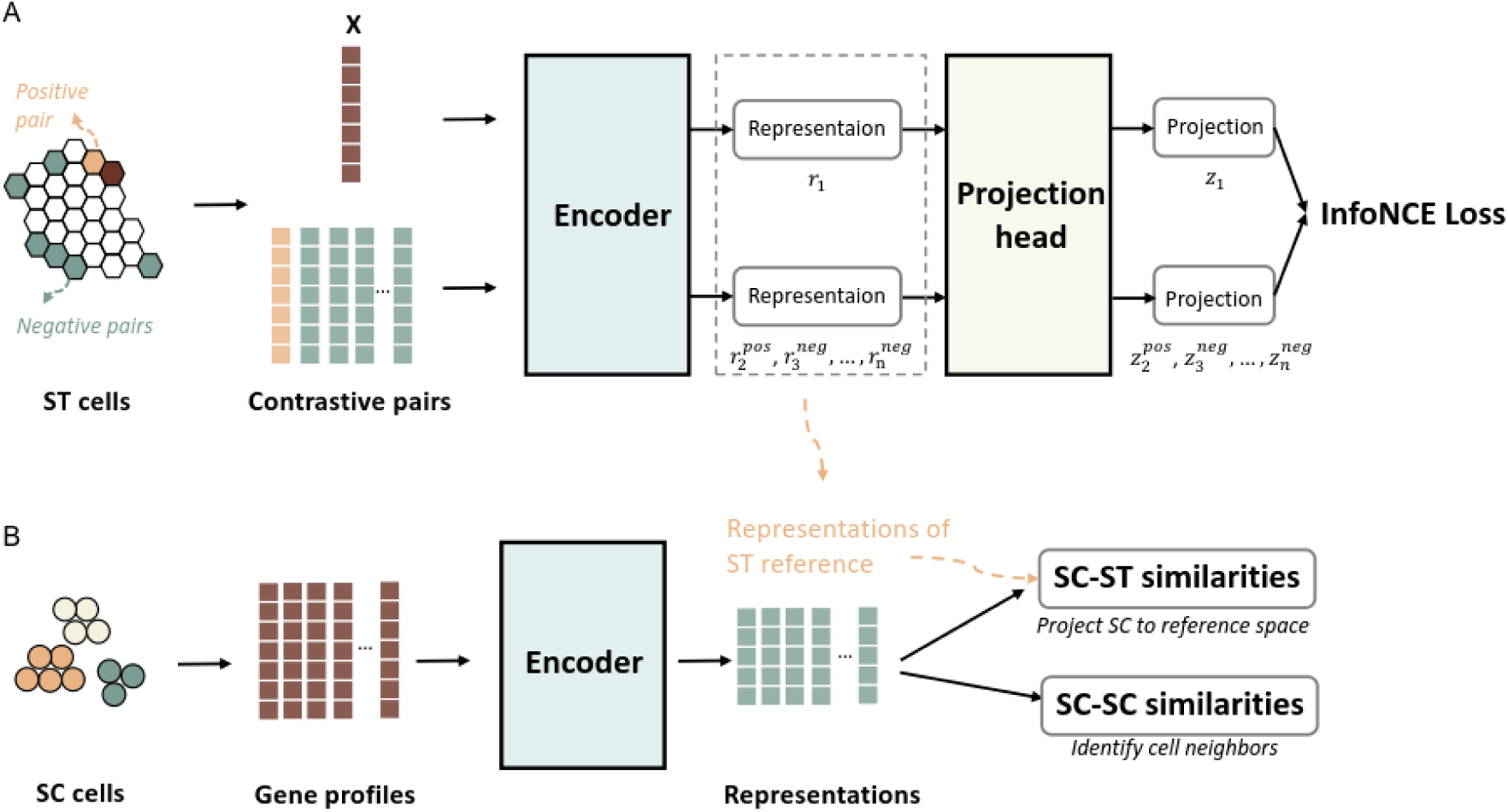
Overview of our method. A. The computational framework. Our method utilized a framework with two key components: an encoder and a projection head. Both components were implemented as MLP. During training, we harnessed spatial transcriptomics (ST) samples as the training dataset. For each sample X, we selected a positive pair from its nearest neighbors and chose *n*-2 random samples as negative pairs. The model’s neural network takes log-normalized gene expressions as input and is optimized through the infoNCE loss, which minimizes the distance between positive pair projections and maximizes the separation of negative pairs. B. Spatial Reconstruction Workflow. The gene profiles of scRNA-seq (SC) samples are processed by the trained encoder to generate representations. These representations enable the identification of cell neighbors and global cell-cell relationships by comparing pairwise SC-SC representation similarities. Moreover, the SC-ST representation similarities was included to locate the most matched ST sample’s coordinates, effectively predicting the SC sample’s spatial location based on the reference dimension.

During inference, the projection head was removed, and the trained encoder received SC gene expressions and generated spatially related representations (Figure 1B). Depending on the downstream tasks, the representations can be used in two ways: 1) obtaining SC-SC similarities to identify cell neighbors and cell-cell distances of the query data, and 2) aligning SC samples to ST samples based on a weighted similarity metric (Methods) and generating coordinates in the reference space.

Evaluation involved two steps. First, we focused on accurately capturing the local cell environment. This was crucial for downstream tasks, such as cell-cell communication and cell-type co-localization. We used two metrics: the average neighbor hit within *k* nearest neighbors, which measures the identification of true individual neighbors; and the Jessen-Shannon distance (JSD) to compare predicted and true neighbor cell-type distributions, considering downstream tasks often rely on detecting the interaction among cell-type groups. Second, we assessed global spatial relationships using the Spearman’s rank correlation coefficient, which compares predicted and true distances among all cells (Methods).

### Benchmarking spatial reconstruction of mouse gastrulation cells using single-cell ST reference

To evaluate the performance of the cell spatial reconstructions of cellContrast, we employed single-cell resolution mouse gastrulation data generated from SeqFISH ^18^. The dataset contains expression profiles of 351 genes collected from three mouse embryos that are on embryonic day (E)8.5 (marked embryos1, 2, and 3). Each mouse embryo has two sections from distinct tissue layers 12μm apart (marked L1 and 2). We trained the model using embryo1 L1 with 10,150 cells, and assessed the performance in two scenarios: 1) testing different layers from the same mouse embryo (embryo1 L2), and 2) testing independent samples (embryo2 L2 and embryo3 L2) (Figure 2A).

**Figure 2.**
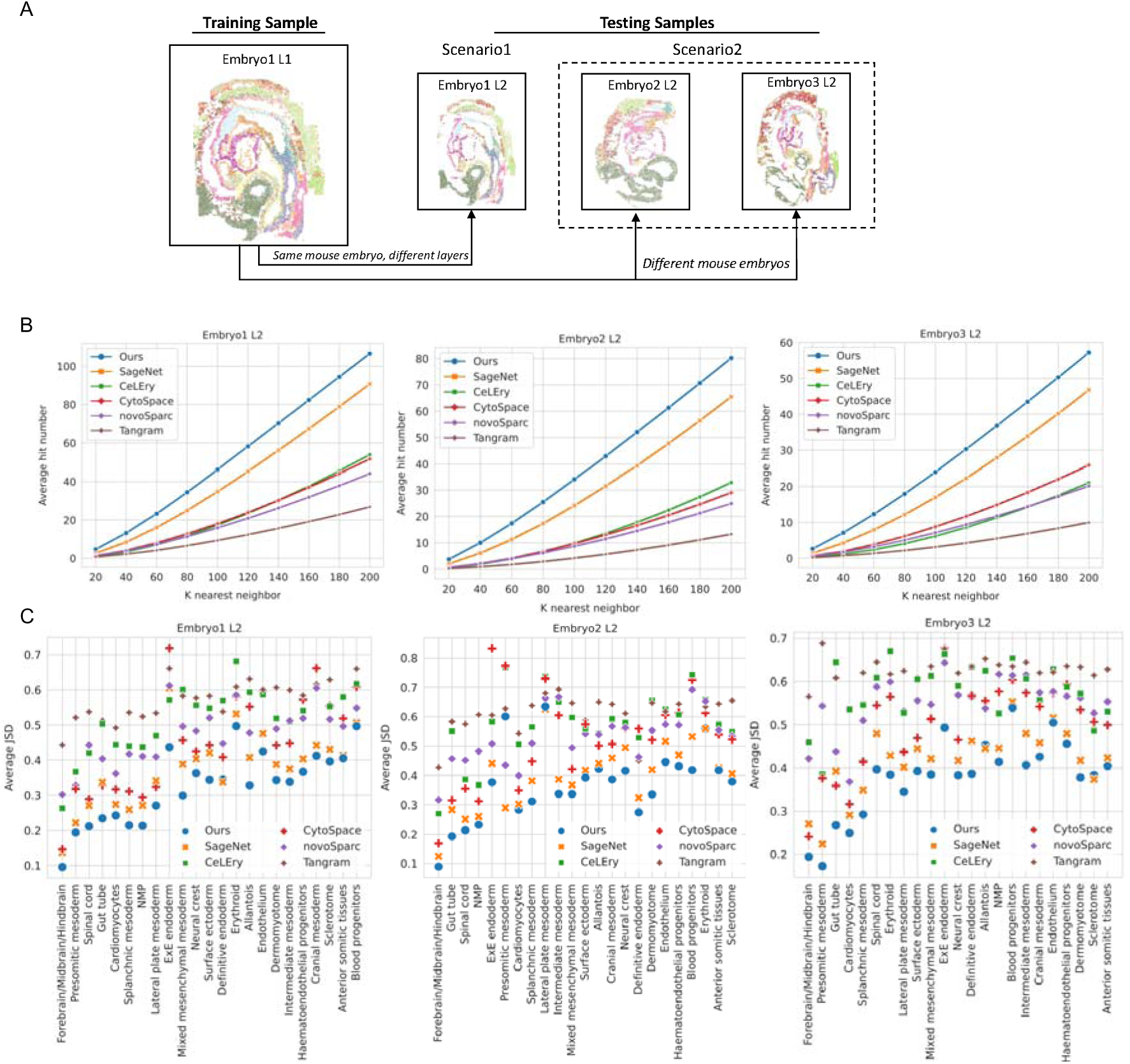
Spatial reconstruction of mouse gastrulation cells using a single-cell resolution ST reference. A. Strategy for evaluating reconstruction performance. The training dataset consistently featured SeqFISH1 L1 data. We investigated two distinct scenarios: 1) evaluation using embryo1 L2 data, sourced from the same embryo as the training data, but originating from different tissues; and 2) assessment using embryo2 L2 and embryo3 L2, derived from independent mouse embryos not utilized in the training phase. B. Evaluation of cell neighbor reconstruction. We quantified the performance of different methods by analyzing the average hit number within various *k* nearest neighbors across the three distinct mouse test datasets. C. Evaluation of neighborhood cell-type reconstruction. We computed the average Jessen-Shannon distance (JSD) for each cell type across the three mouse test datasets.

We conducted a comprehensive comparison of five related methods (Supplementary Data 1-3). Our evaluation encompassed the detection of true cell neighbors within the predicted *k* nearest neighbors surrounding individual cells (Figure 2B). Notably, our method outperformed all others, displaying the highest average hit numbers across all tested datasets. Our method and SageNet managed to identify twice the number of true neighbors than alternative approaches (for example, the median average hit ratios of 47.32% by ours, 36.22% by SageNet and 14.27% by CeLEry). Furthermore, our method outperformed SageNet by 10 additional neighbors when *k* ≥120 (about a 22% gain). Even at lower *k* values, such as 20, our method exhibited an improvement, identifying nearly 1.5 times the number of true neighbors compared to SageNet (4.73 vs 2.69, 3.71 vs 2.01, 2.66 vs 1.43, for L2 of embryo1,2,3, respectively). It is worth noting that all methods yielded a higher average hit number in scenario 1 than in scenario 2. This observation underscores the importance of spatial consistency between training and testing cells, as a stronger alignment enhances the accuracy of predicted neighbors.

We further explored the extent of reconstruction of local cell-type heterogeneity by comparing cell types between the true and predicted nearest 20 neighbors and quantifying their divergence through JSD. A lower JSD value signifies a closer resemblance in cell-type distribution between predicted and true cell neighbors. The results, presented in Table 1, underscore the notable performance of our method, exhibiting the lowest average JSD of 0.237 for embryo1 L2. SageNet was next, attaining a JSD of 0.289, while CytoSpace had a JSD of 0.324. In contrast, other methods displayed average JSDs large than 0.4. Of particular significance, our method maintained consistent JSD performance across independent testing datasets for embryo2 and 3, illustrating its robustness against over-fitting to new contexts. Delving deeper into the JSD averages for individual cell types, as depicted in Figure 2C, our method’s advantage is evident across almost all cell types: for example, gut tube in all three testing samples. The only exception was presomitic mesoderm cells in embryo2 L2, partly due to the larger local cell-type heterogeneity of embryo2 compared to the training embryo1 L1. Worth noting is the discernible trend of increased JSD across all methods as the inherent truth local heterogeneity of cell types escalated.

**Table 1.** Average JSD (*k*=20) and Spearman’s correlation coefficient on SeqFISH1 L1 reference of mouse gastrulation. JSD assesses the consistency of local cell type proportions within k-nearest neighbor. Spearman’s ρ evaluates the correlation of global spatial distances, especially including the non-neighbors as the majority.

| Method | Embryo1 L2 |  | Embryo2 L2 |  | Embryo3 L2 |  |
| --- | --- | --- | --- | --- | --- | --- |
| | JSD | Spearman's $\rho$ | JSD | Spearman's $\rho$ | JSD | Spearman's $\rho$ |
| **Ours** | **0.237** | **0.874** | **0.225** | 0.574 | **0.343** | 0.477 |
| SageNet | 0.289 | 0.543 | 0.273 | 0.514 | 0.391 | 0.391 |
| CeLEry | 0.435 | 0.769 | 0.44 | **0.59** | 0.562 | **0.485** |
| CytoSpace | 0.324 | 0.623 | 0.356 | 0.331 | 0.433 | 0.296 |
| novoSparc | 0.417 | 0.47 | 0.432 | 0.155 | 0.522 | 0.197 |
| Tangram | 0.525 | 0.304 | 0.538 | 0.073 | 0.613 | 0.095 |

\* Spearman's $\rho$ denotes Spearman's correlation coefficient.

The global spatial relationships were assessed by Spearman’s rank correlation between true and predicted pairwise distances for each cell. Our method performs the best in scenario1, while achieved comparable performance with CeLEry in secenario2 (Table 1). Our method, along with SageNet and CeLEry, maintained relatively stable performance across independent mouse embryos, while Tangram, novoSparc, and CytoSpace exhibited significant declines. This divergence in performance may be attributed to the fact that those methods are not focused primarily on mapping SC samples to ST. Although they can perform this task, their main design focuses on ST spot deconvolution based on gene expression, lacking precise spatial arrangement.

### Robustness of spatial location inference across platforms

After validating the efficiency of spatial reconstruction with single-cell reference, we further explored cellContrast in more complicated settings. First, we benchmarked the reconstruction of the cell locations of SC samples with the assistance of a spot-based ST reference. In this experiment, the performance was evaluated in a setting where reference and query cells were: 1) collected from different mouse embryos at slightly mismatched developmental stages; and 2) obtained by different techniques (with StereoSeq and SeqFISH, respectively). The ST reference was spot-based. We took one sample of MOSTA, which was collected from a mouse embryo on E9.5, obtained by Stereo-Seq ^6^. The single-cell resolution embryos1, 2, 3 L2 were included for testing (Figure 3A, Supplementary Data 4-6). There were 5,266 spots with 348 overlapped genes for training the model. As shown in Figure 3B and Supplementary Figure 1, our method achieved the highest mean average hit number of the three test datasets. Specifically, when the nearest neighbor was *k*=200, cellContrast detected 55.388 true cell neighbors. Second-place SageNet got an average hit number of 36.8 and detected 16 fewer than ours. The results suggest the good transferability of our method by capturing complex spatial-dependent features. The reconstructed local cell-type heterogeneity was assessed by JSD for the nearest 20 neighbors. As shown in Table 2, cell neighbors reconstructed by our method identified the most similar cell neighbors with cell-types with the truth with the lowest mean average JSD (0.305 by ours vs. 0.366 by second-best SageNet). The JSDs for each cell type are shown in Figure 3C and Supplementary Figure 2. Our method and SageNet’s kept robust JSDs because they can estimate the query cell-cell distance directly. But methods that can only project the cells into the spatial coordinates based on the ST reference decreased drastically, even for the cell types with low local cell-type heterogeneity (Forebrain/Midbrain/Hindbrain, Gut tube, Cardiomyocytes, etc). Similar patterns were observed for the Spearman’s rank correlation coefficient (Table 2).

**Figure 3.**
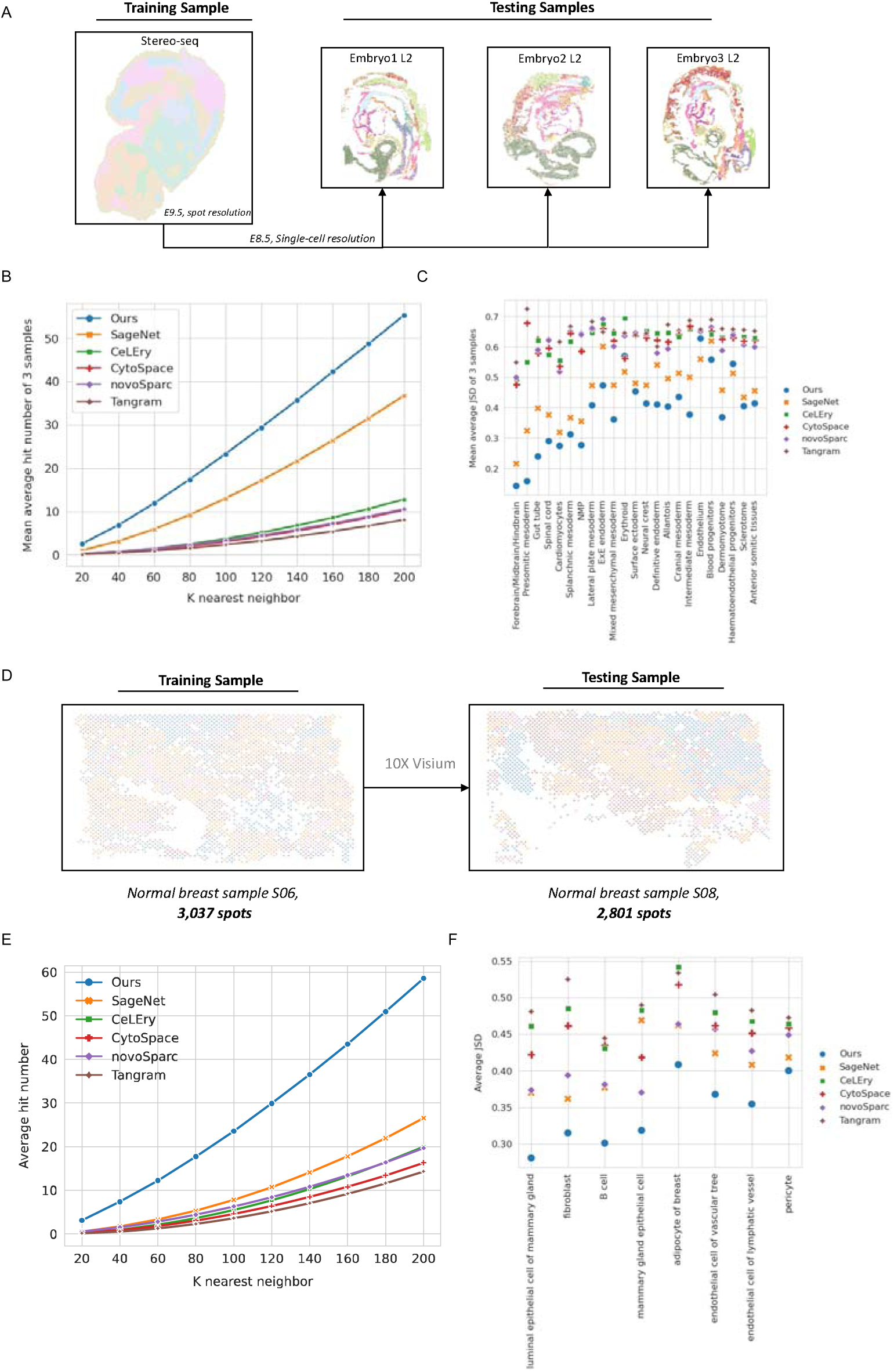
Spatial location inference across platforms. A. Evaluation of spatial reconstruction for the mouse gastrulation cells using array-based ST reference. The training dataset consistently utilized the Stereo-Seq E2S1 sample. The benchmarking was against three independent single-cell resolution ST datasets derived from E8.5 mouse embryos. B. Evaluation of cell neighbor identification. We evaluated the efficacy of different methods by quantifying the average hit number within varying *k* nearest neighbors across the three distinct mouse test datasets; the mean average hit numbers of three testing sets were presented. C. Evaluation of local neighborhood cell-type heterogeneity reconstruction. We computed the average Jessen-Shannon distance (JSD) for each cell type across the three mouse test datasets and showed the mean values of the 3 testing sets. D. Evaluation of spatial reconstruction for 10X Visium human breast cells. The training utilized the normal sample S06, consisting of 3,037 spots, while the testing involved sample S08, comprising 2,801 spots. E. Assessment of cell neighbor identification. We quantified performance by analyzing the average hit number across varying nearest *k* neighbors of the test human breast cells. F. Evaluation of neighborhood cell-type reconstruction of human breast cells using JSD.

**Table 2.** Average JSD (*k*=20) and Spearman’s correlation coefficient of across multiple platform benchmarking experiments.

| Method | Mean of three mouse embryos |  | Human breast |  |
| --- | --- | --- | --- | --- |
| | JSD | Spearman's $\rho$ | JSD | Spearman's $\rho$ |
| **Ours** | **0.305** | 0.283 | **0.306** | 0.103 |
| SageNet | 0.366 | **0.328** | 0.386 | **0.119** |
| CeLEry | 0.577 | 0.156 | 0.465 | 0.077 |
| CytoSpace | 0.568 | 0.099 | 0.444 | 0.005 |
| novoSparc | 0.579 | 0.066 | 0.393 | 0.044 |
| Tangram | 0.614 | 0.053 | 0.485 | 0.007 |

\* Spearman's $\rho$ denotes Spearman's correlation coefficient.

Based on the successful application of our method in spatial reconstruction in mouse embryos with single-cell and spot-based ST reference, we extended our investigation to assess its performance within a recently released spatially resolved genomic atlas of the adult human breast, generated using 10x Visium ^19^. We employed two distinct normal samples – S06 (3,037 spots) and S08 (2,081 spots) – as a training and testing dataset, respectively (Figure 3D, Supplementary Data 7). A total of 36,503 genes were used as input. As SageNet could not handle such a large pool of genes with 1T memory computing server, we used 1,024 highly variable genes as input for it. As a result, our method detected the most true cell neighbors, identifying 58.556 truth neighbors when *k*=200, about two times that of the next-best method (Figure 3E). For reconstructing neighborhood cell-type heterogeneity, our method achieved the most similar neighborhood cell-type distribution of an average of JSD 0.306, followed by SageNet’s 0.386 and NovoSpac’s 0.393; and ours was the most robust for most of the cell types (Figure 3F). Furthermore, in terms of capturing global spatial relationships among cells, the performance among SageNet, ours and CeLEry are comparable.

To further examine cellContrast’s transferability to different techniques, we incorporated the single-cell resolution mouse brain dataset generated by MERSCOPE into our assessment ^20^. Specifically, we utilized cells from the left hemisphere for training and those from the right hemisphere for testing. Similar to our observations with other datasets, our methods demonstrated the highest performance in both local and global spatial reconstruction (Supplementary Figure 3, Supplementary Table 1, Supplementary Data8).

### Unveiling cell-type co-localizations in dissociated mouse gastrulation cells with cellContrast

With the solid foundation laid out in the previous benchmarking sections, we showed the practical efficacy of our method for enhancing biological data analysis. We applied cellContrast to a real-world mouse atlas with embryos at the same developmental stage (E8.5) as the SeqFISH reference [10]. Leveraging the reconstructed spatial coordinates of cells, we showcased the precise detection of cell-type co-localization patterns.

Initially, we built a reference pattern for cell-type co-localizations based on the 6 SeqFISH datasets with truth cell spatial distributions. To identify cell-type spatial associations, we employed Moran’s R from spatialDM, adopting a significance threshold of *p*-value 0.05. Figure 4A presents Moran’s R and *p*-values for all cell-type co-localizations obtained from SeqFISH embryo1 L1. Notably, dataset-specific co-localizations exhibited lower Moran’s R values. “Reference support” for a cell-type co-localization was defined as the number of SeqFISH datasets producing a significant signal for the co-localization pattern. A reference support of 6 denotes the utmost confidence in co-localization, whereas 0 implies the absence of evidence in SeqFISH datasets. Reference supports for each cell pair are shown in Figure 4B. Notably, 88 cell-type co-localizations were identified in the SeqFISH dataset, each with at least one reference support. Intriguingly, a spatial association was detected among blood progenitors, endothelium, and hematoendothelial progenitors, which were closely linked to blood generation ^21^.

**Figure 4.**
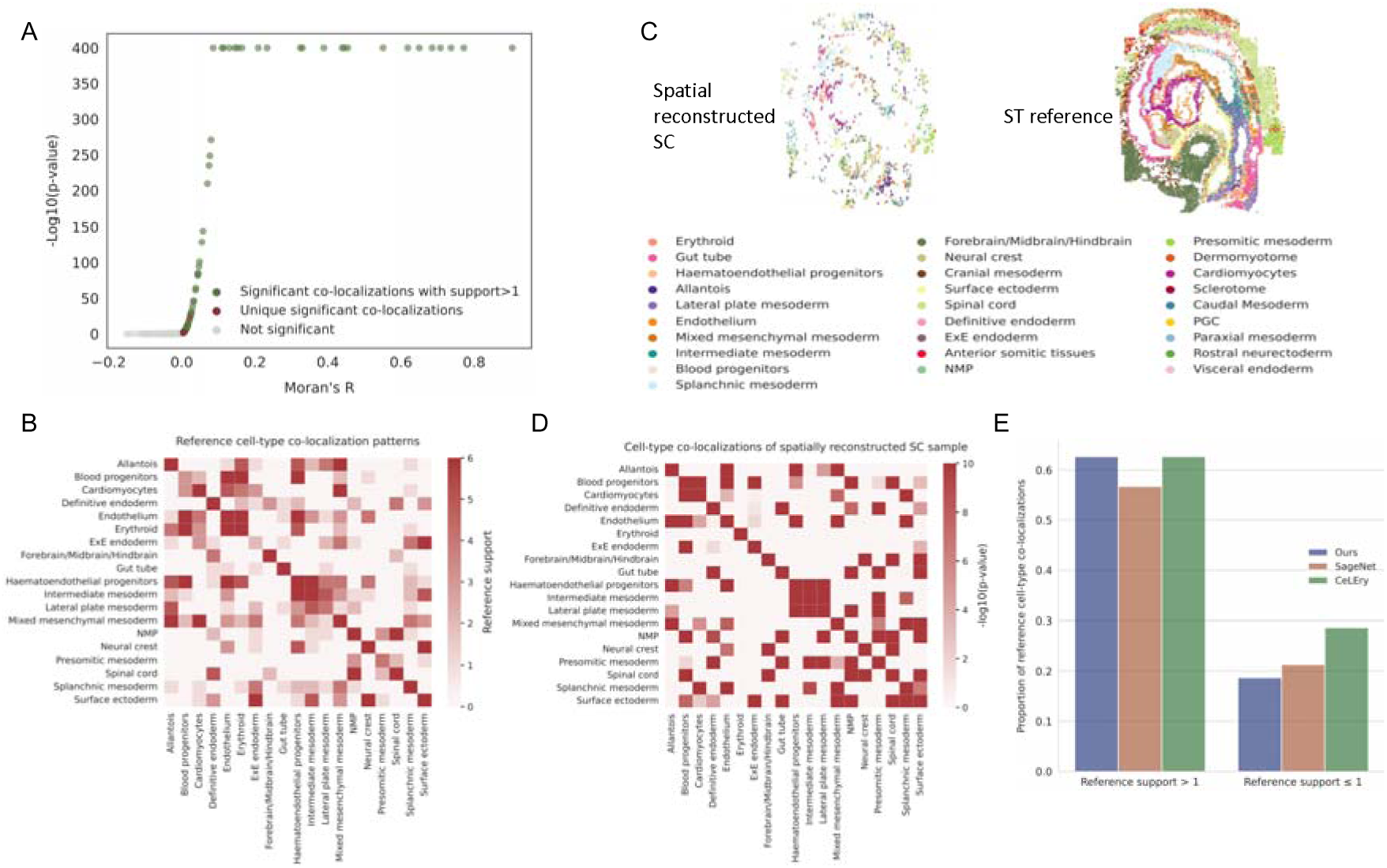
Cell-type co-localization analysis of spatially reconstructed scRNA-seq data of the mouse embryo at E8.5. A. Relationships between Moran’s R and the p-values of all cell-type pairs derived from SeqFISH embryo1 L1. B. Reference cell-type co-localization patterns. We incorporated six SeqFISH datasets; the value of reference support indicates the number of datasets demonstrating significant signals (p ≤ 0.05). C. Reconstructed spatial locations of the SC sample and its ST reference (SeqFISH embryo1 L1). D. Detected *p*-values of cell-type co-localizations based on a spatially reconstructed SC sample. E. Detected co-localization proportions in the SC Sample vs. Reference Patterns.

We further applied our method to a sample of the mouse scRNA-seq atlas. In total, the spatial information of 6,460 cells was reconstructed. The model was trained on SeqFISH embryo1 L1, featuring 350 overlapping genes. The spatial locations of SC cells were achieved by aligning them to the training ST samples. Figure 4C shows the reconstructed spatial locations of SC and reference ST. Employing Moran’s R, cell-type co-localizations were detected, and the corresponding *p*-values are presented in Figure 4D. Importantly, the reference’s major co-localization patterns were captured by spatially reconstructed SC data. We extended our analysis by applying two related spatial reconstruction methods, SageNet and CeLEry, to the same SC sample, and subsequently conducted similar analyses (Methods). We explored detected cell-type pairs with reference support exceeding one. Notably, both our method and CeLEry identified 42 cell-type pairs, while SageNet detected 38 pairs, with 27 co-localizations consistently detected across all three methods. We then employed Fisher’s exact tests for detected and reference co-localizations with support >1. The *p*-values for our method, CeLEry, and SageNet were 2.793e-9, 1.121e-6, and 7.409e-6, respectively (Figure 4E, Supplementary Tables 2, 3, 4). This indicates the effectiveness of spatial reconstruction methods in capturing cell-type co-localization patterns.

### Improved cell-cell communication inference in mouse embryos through reconstructed spatial locations of scRNA-seq data

Cell-cell communication (CCC) analysis benefits from spatial transcriptomic technologies, which provide spatial context and reduce false positives. As shown in Figure 5A, the traditional approach identifies ligand-receptor interactions (LRI) based on gene expressions among different cell groups ^22^. CellChat, a popular tool, uses statistical models to generate the LRIs and the interacting cell groups. With ST, gene expression and spatial locations enable more accurate CCC detection using cell-cell distance constraints. SpatialDM employs Moran’s R to identify co-expression of LRIs and detects interacting local spots or cells. In this study, we demonstrated that our spatial reconstruction method enhances CCC signal accuracy of SC data compared to gene expression alone.

**Figure 5.**
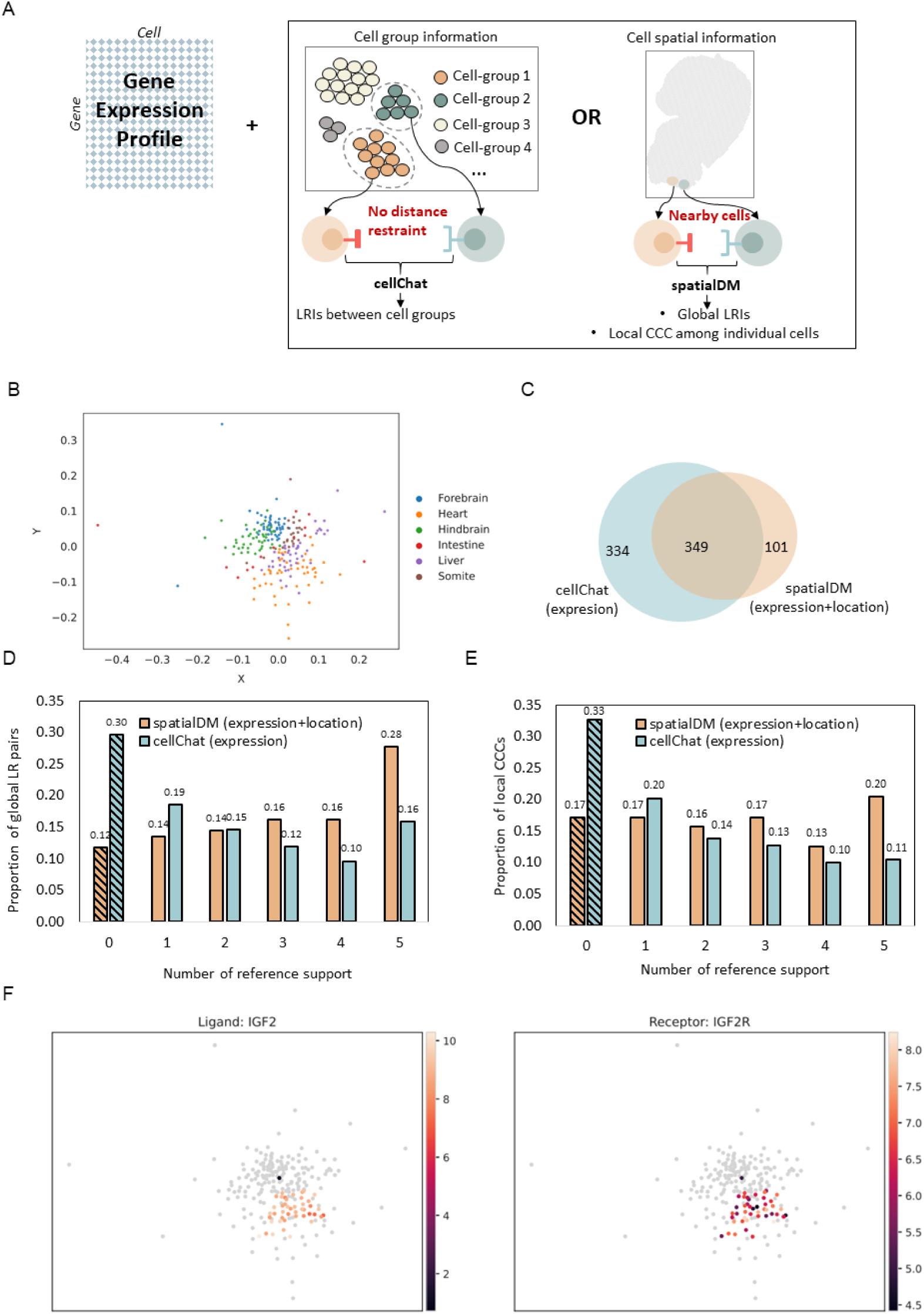
Analysis of cell-cell communication in spatially reconstructed SC data of the mouse embryo at E9.5. A. Demonstration of the cell-cell communication analysis with SC and ST data. We employed CellChat to decipher CCC among different cell groups to compute the co-signals of ligand and receptors based on their gene expression. With the spatial distance constraint incorporated by ST data, SpatialDM detected the global LRIs and the local CCCs among individual cells. B. Reconstructed locations of E1. The spatially reconstructed locations of E1 were accomplished by leveraging the predicted cell-cell dissimilarities into a 2D space through the MDS algorithm. C. Comparison of global ligand-receptor pairs. We compared the identification of global ligand-receptor pairs using cellChat and spatialDM. Notably, cellChat exclusively employs E1’s gene expression data, while spatialDM integrates both gene expression and predicted spatial cell locations. D. Reference support for global ligand-receptor pairs. E. Reference support for local cell-cell communication. F. Spatial demonstration of the detected ligand-receptor interaction of LGF2-LGF2R.

We reconstructed the spatial locations of mouse embryo scRNA-seq sample S1 (scRNA-S1), derived from GES87038. The sample was collected on E9.5 with cells from forebrain, hindbrain, heart, liver, intestine, and somite tissues ^23^. The model was trained with one Stereo-Seq sample, using 444 genes that are highly variable and overlapped with the scRNA-S1. The pair-wise cell-cell similarities were derived from the learnt representations of the cells and can be leveraged to 2D space using the multidimensional scaling (MDS) algorithm (Figure 5B). Notably, forebrain and hindbrain cells are clearly grouped into different clusters and near each other, suggesting cell-cell spatial relationships were well characterized.

Subsequently, we conducted two distinct analyses for cell-cell communication (CCC) detection. Initially, we employed cellChat to detect ligand-receptor interactions (LRIs) based solely on gene expression across various tissues. Then we employed spatialDM to identify LRIs and associated interacting cells by incorporating the reconstructed cell positions derived from the scRNA-S1 (Methods). First, we examined global LRIs, by ignoring specific interacting tissues or cells. Both methods identified a total of 349 global LR pairs, with cellChat detecting more unique LRIs (Figure 5C). To assess the reliability of these LRIs, we established reference CCCs through the application of spatialDM on five Stereo-seq samples possessing ground-truth cell locations. These samples originated from two mouse embryos at developmental stage E9.5, aligning temporally with the scRNA-S1. The reference support for an LRI was defined as the number of Stereo-seq datasets that produced the same signals. A higher reference support indicated enhanced reliability in the LRI. As shown in Figure 5D, where spatialDM consistently identified a greater portion of LRIs with reference support > 1 (for example, 0.28 VS 0.16 for 5 reference support), accompanied by a reduction in LRIs with no support (0.12 VS 0.30). In contrast, for a large proportion of cellChat identified LRIs, no evidence has been found in the reference datasets. This result underscores the enhanced precision of LRI detection facilitated by the incorporation of reconstructed cell spatial information.

In addition to global LRIs, we further explored local CCCs, which are vital for understanding the role of the cells in the local context. We focused on tissues that overlapped the reference datasets – forebrain, hindbrain, heart, and liver – for validation. We evaluated local CCCs using “Local spot evidence” to denote matched spots inferred from reference datasets, while “Number of reference support” denotes the number of reference datasets that have at least one local spot evidence for the corresponding CCC (Methods). We obtained similar conclusions to those of global LRIs (Figure 5E): the reconstructed cell locations notably reduced local unsupported CCCs, minimizing potential false positives.

As an example, we examined the established insulin-like growth factor 2 (IGF2) and its high-affinity receptor IGF2R, which is well known for trophoblast morphogenesis regulation [13]. Forty-five cells within the scRNA-S1, primarily from heart and liver tissues, exhibited significant IGF2-IGF2R signals using spatialDM. The normalized expressions and reconstructed spatial coordinates of these cells are shown in Figure 5F, effectively supporting cell-cell communication within a short range of reconstructed physical distances.

## Discussion

In this study, we introduced a novel method, named cellContrast, designed to reconstruct spatial relationships among SC based on gene expression profiles, a task of paramount importance in deciphering complex biological activities. CellContrast leverages a deep contrastive learning framework with an encoder-projector structure. This design enables the projection of gene expressions into a latent space, where physically proximate cells exhibit similar representation values. Achieving this involves constructing training batches containing contrastive pairs, derived from ST data with truth locations and optimizing them using the infoNCE loss. The resulting cell-cell distances are estimated through cosine similarities of learned representations, which can be mapped to location coordinates using ST reference or MDS algorithms.

To evaluate cellContrast, we conducted extensive benchmarking experiments on diverse datasets, including mouse embryo, human breast and mouse brain samples. These datasets included various platforms, ranging from spot-based (Stereo-Seq and 10X Visium) to single-cell-based resolution STs (SeqFISH, MERSCOPE). Our method’s performance and robustness were assessed in different scenarios: 1) training and testing on distinct layers of the same mouse embryo with single-cell resolution; 2) training and testing on different mouse embryos, both at single-cell resolution; 3) training on spot-based resolution data and testing on single-cell resolution data from different mouse embryos; and 4) both training and testing on spot-based resolution data with independent human breast samples. 5) training and testing on single-cell resolution MERSCOPE data with mouse brain cells. In all benchmarking scenarios, cellContrast consistently outperformed existing methods in local spatial reconstruction, such as SageNet ^14^, CeLEry ^15^, CytoSpace ^13^, novoSpaRc ^11^, and Tangram ^12^, demonstrating its effectiveness and transferability in capturing local spatial relationships among neighboring cells. Our contrastive model primarily focuses on learning features related to neighboring cells, treating all distant cells equally. To address the challenge of estimating global spatial distance relationships, we project the SC samples onto reference ST locations based on the similarities of the learnt representations. This projection employs a weighted similarity metric based on both SC neighbors and ST samples. Notably, when evaluating global reconstruction performance using Spearman’s rank correlation coefficient, we observed improved results by incorporating more predicted nearest SC neighbors (as shown in Supplementary Table 5). However, this approach also introduces a potential issue known as “cell clumping,” where multiple neighbors are projected into the same location. The trade-off between accurately ranking global distances and addressing cell clumping can be adjusted by the number of SC neighbors used for SC-ST alignment. On the other hand, we believe that in the task of spatial reconstruction, more emphasis should be placed on the local environment. This emphasis is rooted in the fact that neighboring cells are often actively engaged in mutual interactions, such as paracrine and juxtacrine signaling. Conversely, non-neighboring cells function within a more complex system, often requiring the involvement of mediators, such as blood plasma ^22^, where the importance of spatial reconstruction diminishes. Additionally, contemporary downstream biological algorithms typically consider the interactions among neighboring cells in their calculations.

Reconstructing the spatial locations of SC data unlocked the potential for two crucial downstream analyses, exemplified with real-world SC samples in this study. The first is cell-type co-localizations analysis: using cellContrast, we reconstructed spatial locations for cells from a mouse embryo on E8.5 and detected cell-type co-localization patterns with Moran’s R. To validate our findings, we built reference patterns from six ST datasets with truth cell locations. Remarkably, the detected co-localizations aligned well with the reference patterns, exhibiting substantial agreement across the three spatial reconstruction methods (cellContrast, SageNet, and CeLEry). This analysis demonstrated that spatially reconstructed SC data can effectively capture cell-type co-localization patterns, thus enhancing biological insights. The second is cell-cell communication (CCC) analysis, which is crucial for comprehending cell differentiation and tissue organization. While many methods infer CCCs from SC data, the lack of spatial constraints might introduce false positives. We employed cellContrast for CCC analysis on E9.5 mouse embryo SC data and compared CCCs inferred solely from gene expression (with cellChat) to those incorporating spatial information (with spatialDM). Validated against five ST datasets, our results exhibited reduced false positives in CCCs when spatial information was considered. We also noted that because the reference CCCs were identified through the same approach with the spatial reconstructed SC data, the results could be advanced to it to some extent. However, we consider the overall effect of spatial reconstruction to remain the same.

In contrast to ongoing efforts to enhance ST data resolution and impute missing genes through SC reference, our study adopts an inverse approach. Leveraging the wealth of SC data available, which covers a broader range of samples and species than available ST data, we present a novel solution for spatial relationship inference. This approach allows us to capitalize on the spatial patterns embedded in ST data to reconstruct the spatial relationships of SC samples, when they were collected from the same region of the same species. A certain level of mismatch may exist between the query SC data and the ST reference due to limited ST resources (e.g. the developmental stages of embryos), but our method shows robustness, thanks to its primary aim of learning from a spatially preserved representation instead of explicit translation of spatial location.

Looking ahead, cellContrast holds significant promise for comprehensive utilization of SC data in diverse contexts. Future directions include refining the method’s performance with improved ST data and exploring novel applications of spatial reconstruction in the realm of single-cell analysis.

In conclusion, our proposed method, cellContrast, offers a powerful solution for accurate spatial relationship reconstruction in SC data. By harnessing the intricate spatial patterns inherent in ST data, cellContrast provides a tool for enhancing various downstream analyses and facilitating critical insights into cellular behavior. It provides the way for deeper exploration of spatial dynamics in single-cell research and offers opportunities for new discoveries reliant on spatial information.

## Methods

### Implementation of cellContrast

#### Training batch construction

Constructing positive and negative cell pairs within training batches is a crucial aspect of contrastive learning. The primary goal of the cellContrast is to project cells into a latent space where spatially proximate cell pairs are similar. This requires careful formation of training batches, where neighboring cells are considered as positive pairs, and non-neighboring cells act as negative pairs.

Given a batch comprising *N* cells, initially, *N*/2 cells are randomly chosen from the training dataset. For each cell X from the *N*/2 pool, one of the *k*-nearest spatial located neighbors to X is considered to be a positive pair. The neighboring size *k* is determined based on the specific dataset: for the SeqFISH, MERSCOPE, Stereo-Seq, Visium 10X training data, values of 80, 80, 20, and 20, respectively, were used. This adjustment takes into account the decreased spatial resolution associated with the above four Spatial ST technologies. Besides the positive cell pairs in the training batch, the remaining cell pairs are treated as negative pairs. The batch size *N* was set as 64 in this study.

#### Model structure and training

CellContrast is composed of an encoder and a projection head, both implemented as MLP. In this study, there were two layers for both the encoder and the projection head. The node numbers were (1024, 512) for the encoder and (256, 128) for the projection head. For the encoder, BatchNorm was added for each layer and leaky ReLU was used as the activation function. For the projection head, ReLU was added to the first layer of MLP.

In the training phase, cellContrast accepted log-normalized gene expressions, which were calculated by scran ^24^. The projections were employed to calculate the infoNCE loss (Eq.1).

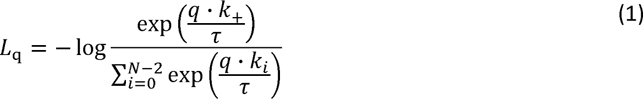

Where *L*_q_ denotes the loss of sample *q*. *τ* means the temperature parameter, which was set as 0.05 in this study. *k*_+_ denotes the positive pair of sample *q*, and *k*_i_ denotes the *i*^th^ the negative sample. *N* denotes the batch size, and *N*-2 is the number of negative samples.

The loss of each training sample was obtained by the above, and the batch mean loss value was used for subsequent optimization. The optimizer of SGD (momentum=0.9, weight_decay=5e-4) was adopted. The learning rate of the initial value 0.1 was scheduled with cosine decay. The model was trained 3000 epochs.

#### Spatial location inference of SC data

During the inference phase, the projection head was discarded, because the encoder is better on learning positive pairs’ spatial invariance ^16, 25^. and log-normalized gene expressions were fed into the previously trained encoder to generate representations. The cosine similarities among representations were utilized to predict spatial distances. Two different approaches were employed. First, to identify the neighbors among the query SC samples, SC-SC similarities were calculated. The top *k* samples with the highest cosine similarities were regarded as the nearest neighbors. Additionally, the coordinates of the SCs could be projected onto a pseudo space using the MDS algorithm based on the pairwise SC-SC similarities. The second approach is aligning SC to ST samples. This alignment is achieved through a weighted similarity metric that takes both SC-SC and SC-ST similarities into account. For a given query SC sample, we aim to estimate its spatial similarity to all available ST samples. This spatial similarity is represented as **S**:

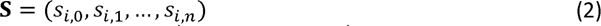

where *s_i,j_* denotes the spatial similarity between the *i*^th^ SC sample and *j*^th^ ST sample, with *N* being the total number of reference ST samples. And *s_i,j_* is calculated based on the following:

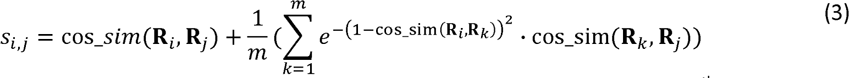

where cos_sim represents the cosine similarity, **R***_i_* is the learnt representations of *i*^th^ SC sample and **R***_k_* is the representation of predicted *k*^th^ nearest SC neighbor of *i*^th^ SC sample. **R***_j_* denotes the learnt representation of *j*^th^ ST sample. M denotes the nearest *m* SC neighbors inferred from SC-SC similarities. This procedure is equivalent to smoothing the original SC-ST distance by the inferred SC-SC neighborhood graph. By default, M is set as 1 (no SC neighbor was included) in the benchmarking experiments.

Subsequently, we align each SC sample with the ST sample that demonstrates the maximum weighted similarity. The coordinates of this matched ST sample are considered as the predicted location for the SC sample.

For downstream tasks that require cell coordinates as input, such as cell-type co-localization and cell-cell communication, we utilized the first approach for the reference of spot-level resolution, and the second approach for the reference of single-cell resolution.

### Benchmarking analysis with real-world datasets

#### Related methods

To fully assess cellContrast, we performed benchmarking experiments with several related methods, including sageNet ^14^, CytoSpace ^13^, Tangram ^12^, novoSparc ^11^, and CeLEry ^15^. The running and evaluation scripts of the benchmarking analysis can be found at: https://github.com/HKU-BAL/CellContrast

##### SageNet

SageNet employed a graph neural network to reconstruct the latent spatial locations by modeling the interactions between cells and genes. The model generated predicted pairwise cell-cell distance for the query samples. For the seqFISH training data, we partitioned the data and set the regularization parameters of the graphical lasso (GLASSO) to be the same as those reported by the authors for the same dataset. In the case of training with the Stereo-Seq and 10X Visium samples, which are both array-based ST technologies, we created three distinct partitions by assigning the spots into 2x2, 3x3, and 4x4 square grids and set the regularization parameters to 0.5 and 0.75, respectively, in line with the authors’ approach for spot-based spatial datasets. All models were trained for 20 epochs using the default settings. For the benchmarking experiment of the human breast, 1024 highly variable genes were used for training, as our 1TB memory computing server was not capable of handling all available genes in the dataset. All genes were used in other experiments.

##### CytoSpace

CytoSpace assigns SC samples to spatial locations by minimizing a correlation-based cost function for gene expressions between query and reference data. Thus, it returns the most closely matched reference spatial coordinates for the query SC samples. In our benchmarking experiments, the neighbors and cell-cell distances of each query sample were determined using the Euclidean distance of the predicted coordinates. It should be noted that CytoSpace may not be capable of allocating positions to all samples, and the assigned positions could not be unique, so the evaluation of CytoSpace was limited to successfully assigned samples, and we adopted the first appearance of predictions. In benchmarking experiments that use single-cell resolution seqFISH samples as the spatial reference, we configured CytoSpace with the option “-single-cell” and supplied the author annotated cell types for both the reference and query datasets. The parameter “-noss” was set at 10000. For experiments that use spot-based Stereo-Seq and 10X Visium samples as the spatial reference, we ran CytoSpace with its default parameters.

##### Tangram

Tangram is similar to CytoSpace in that it seeks to learn mapping functions from SC cells to ST samples through deep learning. At the cell level, Tangram generates the probability of SC samples being present in all available reference ST positions. In our benchmarking experiments, we selected the reference position with the maximum probability as the predicted spatial location. We ran Tangram with all genes and default parameters for all benchmarking settings.

##### novoSpaRc

NovoSpaRc is a computational method that assigns cells to tissue locations by assuming that cells with similar gene expressions are closer in spatial locations. Like Tangram, NovoSpaRc produces mapping probabilities for SC samples to locations of the reference. Therefore, we selected the positions with the maximum probability as the prediction in our benchmarking experiments. The predefined locations were constructed based on the reference ST dataset, which were as indicated in Figures 2A, 3A, and D. Following the NovoSpaRc analysis pipeline, we first identified marker genes from the reference dataset and then performed the mapping process based on these markers. As per the authors’ reported practice, we set num_neighbors_s and num_neighbors_t as 5, alpha_linear as 0.8, and epsilon as 5e-3.

##### CeLEry

CeLEry is a supervised deep-learning model designed to recover the spatial domain and locations of SC datasets. It achieves this by reconstructing the individual cell’s spatial location and producing spatial coordinates. We trained CeLEry with the default parameters for all benchmarking experiments and inferred the predicted cell neighbors and cell-cell distances by calculating the Euclidean distance of the predicted coordinates.

#### Cell-type co-localization analysis

We employed SpatialDM’s Moran’s R ^26^ to investigate the cell-type co-localization patterns of ST data and spatially reconstructed SC data. To utilize it, we encoded a binary variable (0 or 1) for each location to signify whether or not it is occupied by a specific cell type. The significant spatial associations between different cell types were determined based on *p*-value ≤ 0.05.

To construct reference patterns, we analyzed 6 SeqFISH datasets from 3 mouse embryos. We set *n_neighbors* to 20 and the radial basis kernel (*l*) to 0.1 to identify the reference patterns. For the spatial reconstructed SC sample of cellContrast, we set the *l* as 0.1. And for ceLEry’s, we set the *l* as 0.1 and *n_neighbors* as 20. While SageNet does not directly output the coordinates of SC samples, we used MDS to project its produced cell-cell distance as pseudo space and set the *l* as 1e-4, and *n_neighbors* as 20, because the scale of predicted distance is very small. The parameters of spatialDM were tuned based on the visualization of the *weight_matrix* according to spatialDM guidance (Supplementary Figure 4).

#### Cell-cell communication analysis

In the cell-cell communication analysis, we employed two methods. We used CellChat to detect LRI among different tissues of the SC mouse embryo data using gene expression only [13]. We used the recommended analysis pipeline and set the parameter “type” as “truncatedMean”. To detect CCCs in the spatially reconstructed SC data and reference ST data, we used spatialDM [12]. We set the parameter “*n_neighbors*” as 20, and the “*l*” as 0.1 for all. We set the other parameters to default. We kept only the results with p-value ≤ 0.05.

To validate the detected CCCs, we compared the detected global LRI and local CCCs with the 6 reference ST datasets. For global LRIs, we defined “reference support” as the number of datasets that detected at least one same LRI, regardless of where it was detected. We defined local CCCs as a combination of an LRI and where it was detected. The “local spot evidence” was defined as follows: for identified CCCs of CellChat, which are a combination of a specific LRI (X) and the involved tissue groups (A and B), local spot evidence refers to at least one spot with significant X in A and B detected in the individual reference dataset. For identified CCCs of spatialDM, which were a combination of LRI X and the specific cell C, the local spot evidence was defined if the same X was detected in the spots in the tissues where C was located in the individual reference ST dataset. The reference support denotes the number of datasets that have local spot evidence. Only results found in tissues that overlapped with the reference were included in the analysis of local CCCs.

#### Evaluation metrics

We concluded that the performance of spatial reconstruction of SC samples could be evaluated by the following criteria: 1) how many truth neighbors can be found; 2) how well the local cell-type heterogeneity can be preserved, which is fundamental for understanding many biological processes; and 3) whether global cell-cell relationships can be captured. To achieve these goals, we used three evaluation metrics: nearest neighbor hit, Jessen-Shannon distance, and the Spearman’s rank correlation.

The nearest neighbor hit refers to the number of successfully detected true neighbors within the *k* nearest neighbors. This is the strictest metric that requires individual-level spatial reconstruction correctness. For each test sample, we can obtain its corresponding nearest neighbor hit, and the average nearest neighbor hit denotes the mean values for all testing samples. In this study, we tested the performance by varying *k* values of 20, 40, …, 200.

To evaluate local cell-type heterogeneity, we adopted the Jessen-Shannon distance (JSD) to compare the cell-type distributions of predicted *k* nearest neighbor cells and the true *k* nearest neighbors. K was set as 20. The ratios for each cell type of the predicted neighbors and truth neighbors were obtained and then JSD was calculated using the jensenshannon function in the Scipy package. Further, the average JSD was calculated for all test samples, as well as by cell-type groups.

To evaluate global spatial reconstruction performance, we employed the Spearman’s rank correlation coefficient. Given a test sample, we obtained its coordinates by SC-ST mapping approach and the pairwise Euclidean distances between it and all other test samples. Then, the truth distances can be obtained through true locations. We incorporated the Spearman’s rank correlation to assess the monotonic relationship between the predicted and true distances. We calculated the average Spearman’s rank correlation coefficient across all query cells as a measure of the global spatial reconstruction performance.

## Data availability

The original data used in this study can be obtained at the following links: (1) seqFISH data on mouse organogenesis [https://crukci.shinyapps.io/SpatialMouseAtlas/]; (2) Stereo-seq data on mouse embryo [https://db.cngb.org/stomics/mosta/download/, E9.5_E2S1]; (3) 10X Visium human breast data [https://cellxgene.cziscience.com/collections/4195ab4c-20bd-4cd3-8b3d-65601277e731, S06, S08]; (4) MERSCOPE mouse brain data [https://info.vizgen.com/mouse-brain-data, s1r1] (5) scRNA-seq data of mouse embryo on E9.5 obtained from the GEO database (GSE87038); and (6) scRNA-seq data of mouse embryo on E8.5 [https://content.cruk.cam.ac.uk/jmlab/atlas_data.tar.gz, s29].

## Code availability

https://github.com/HKU-BAL/CellContrast

## Authors’ contributions

RL and SL conceived the project. YH supervised the study. SL and JM designed the algorithm. SL implemented the package and performed all data analysis, with support from YH and RL. TZ, YJ and BL participated the benchmarking. YH, RL, SL, JM involved in drafting the manuscript. All authors read and approved the final manuscript.

## Supporting information

Description of Supplementary Data

## References

1. Williams, C.G., Lee, H.J., Asatsuma, T., Vento-Tormo, R. & Haque, A. An introduction to spatial transcriptomics for biomedical research. Genome Medicine 14, 68 (2022).

2. Eng, C.-H.L. et al. Transcriptome-scale super-resolved imaging in tissues by RNA seqFISH+. Nature 568, 235–239 (2019).

3. Wang, X. et al. Three-dimensional intact-tissue sequencing of single-cell transcriptional states. Science 361, eaat5691 (2018).

4. Chen, K.H., Boettiger, A.N., Moffitt, J.R., Wang, S. & Zhuang, X. RNA imaging. Spatially resolved, highly multiplexed RNA profiling in single cells. Science 348, aaa6090 (2015).

5. Rodriques, S.G. et al. Slide-seq: A scalable technology for measuring genome-wide expression at high spatial resolution. Science 363, 1463–1467 (2019).

6. Chen, A. et al. Spatiotemporal transcriptomic atlas of mouse organogenesis using DNA nanoball-patterned arrays. Cell 185, 1777–1792.e1721 (2022).

7. Kleshchevnikov, V. et al. Cell2location maps fine-grained cell types in spatial transcriptomics. Nature biotechnology 40, 661–671 (2022).

8. Cable, D.M. et al. Robust decomposition of cell type mixtures in spatial transcriptomics. Nature biotechnology 40, 517–526 (2022).

9. Abdelaal, T., Mourragui, S., Mahfouz, A. & Reinders, M.J.T. SpaGE: Spatial Gene Enhancement using scRNA-seq. Nucleic acids research 48, e107–e107 (2020).

10. Qiao, C. & Huang, Y. Reliable imputation of spatial transcriptome with uncertainty estimation and spatial regularization. bioRxiv, 2023.2001.2020.524992 (2023).

11. Moriel, N. et al. NovoSpaRc: flexible spatial reconstruction of single-cell gene expression with optimal transport. Nat Protoc 16, 4177–4200 (2021).

12. Biancalani, T. et al. Deep learning and alignment of spatially resolved single-cell transcriptomes with Tangram. Nature methods 18, 1352–1362 (2021).

13. Vahid, M.R. et al. High-resolution alignment of single-cell and spatial transcriptomes with CytoSPACE. Nature biotechnology (2023).

14. Elyas, H. et al. Supervised spatial inference of dissociated single-cell data with SageNet. bioRxiv, 2022.2004.2014.488419 (2022).

15. Zhang, Q. et al. Leveraging spatial transcriptomics data to recover cell locations in single-cell RNA-seq with CeLEry. Nature communications 14, 4050 (2023).

16. Chen, T., Kornblith, S., Norouzi, M. & Hinton, G. in International conference on machine learning 1597-1607 (PMLR, 2020).

17. Oord, A.v.d., Li, Y. & Vinyals, O. Representation learning with contrastive predictive coding. arXiv preprint arXiv:1807.03748 (2018).

18. Lohoff, T. et al. Integration of spatial and single-cell transcriptomic data elucidates mouse organogenesis. Nature biotechnology 40, 74–85 (2022).

19. Kumar, T. et al. A spatially resolved single-cell genomic atlas of the adult human breast. Nature 620, 181–191 (2023).

20. Vizgen (https://info.vizgen.com/mouse-brain-data; 2022).

21. Pinheiro, I. et al. Discovery of a new path for red blood cell generation in the mouse embryo. Experimental Hematology 44, S94 (2016).

22. Armingol, E., Officer, A., Harismendy, O. & Lewis, N.E. Deciphering cell–cell interactions and communication from gene expression. Nature Reviews Genetics 22, 71–88 (2021).

23. Dong, J. et al. Single-cell RNA-seq analysis unveils a prevalent epithelial/mesenchymal hybrid state during mouse organogenesis. Genome biology 19, 31 (2018).

24. Lun, A.T., McCarthy, D.J. & Marioni, J.C. A step-by-step workflow for low-level analysis of single-cell RNA-seq data with Bioconductor. F1000Res 5, 2122 (2016).

25. Ma, J., Hu, T. & Wang, W. Deciphering the Projection Head: Representation Evaluation Self-supervised Learning. arXiv preprint arXiv:2301.12189 (2023).

26. Li, Z., Wang, T., Liu, P. & Huang, Y. SpatialDM for rapid identification of spatially co-expressed ligand–receptor and revealing cell–cell communication patterns. Nature communications 14, 3995 (2023).

