## Supplementary Information for "CellContrast: Reconstructing Spatial Relationships in Single-Cell RNA Sequencing Data via Deep Contrastive Learning"

### 1. Supplementary Figures

A

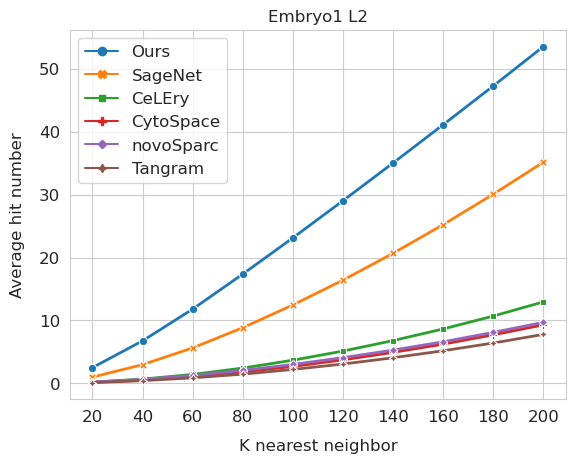

B

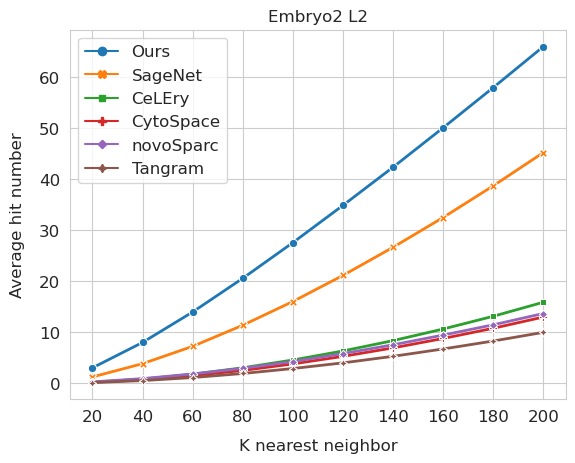

C

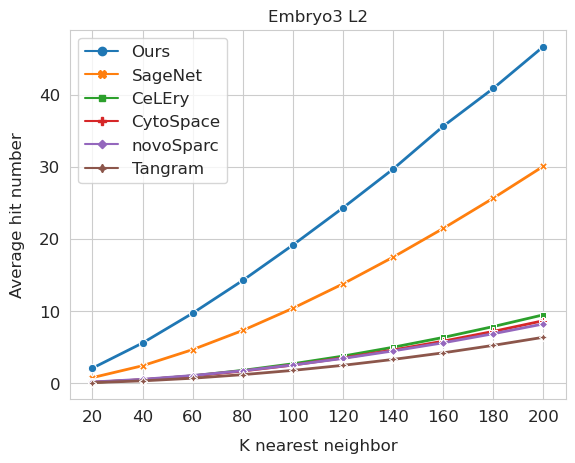

Supplementary Figure 1. Evaluation of neighbor reconstruction in mouse gastrulation cells using array-based ST reference. A. Average hit number with varying *k* nearest neighbors for testing dataset embryo1 L2. B. Average hit number with varying *k* nearest neighbors for testing dataset embryo2 L2. C. Average hit number with varying *k* nearest neighbors for testing dataset embryo3 L2.

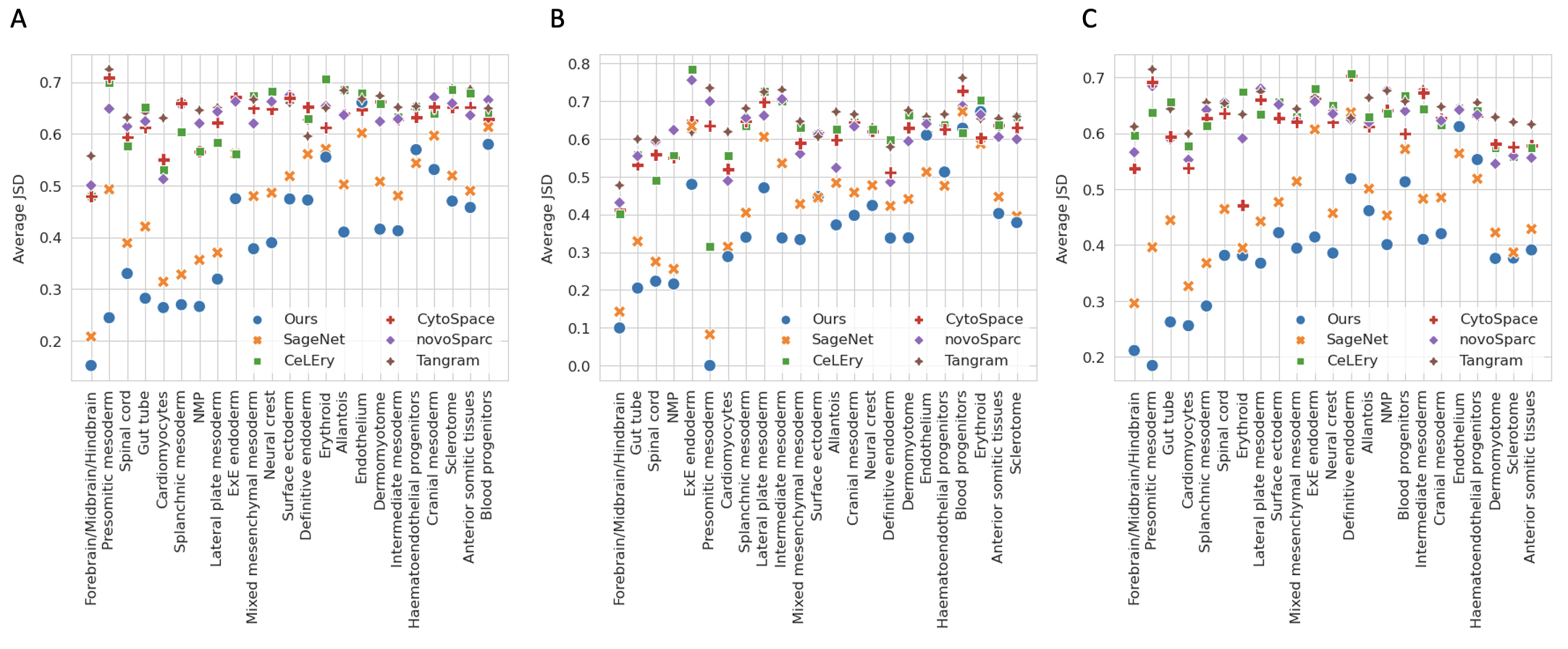

Supplementary Figure 2. Evaluation of local neighborhood cell-type heterogeneity in mouse gastrulation cells using array-based ST reference. A, Jessen-Shannon distance of cell types for testing dataset embryo1 L2. B, Jessen-Shannon distance of cell types for testing dataset embryo2 L2. C, Jessen-Shannon distance of cell types for testing dataset embryo3 L2

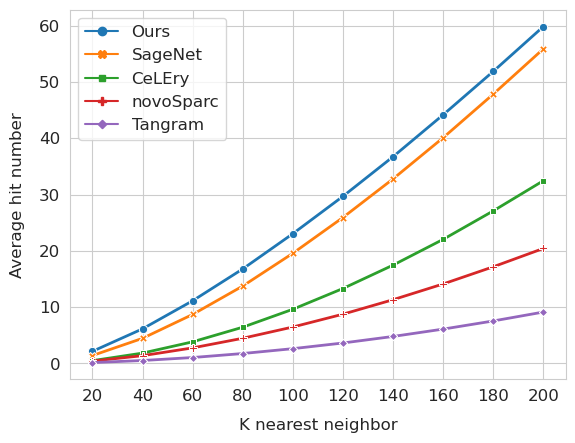

Supplementary Figure 3. Evaluation of neighbor reconstruction in mouse brain cells generated by MERCOPE. Noted that CytoSpace was excluded from the analysis due to the absence of cell type annotations, as cell type information is a required parameter for its single-cell mode.

A

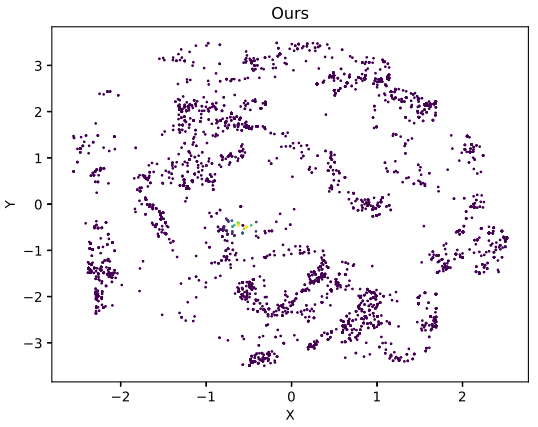

B

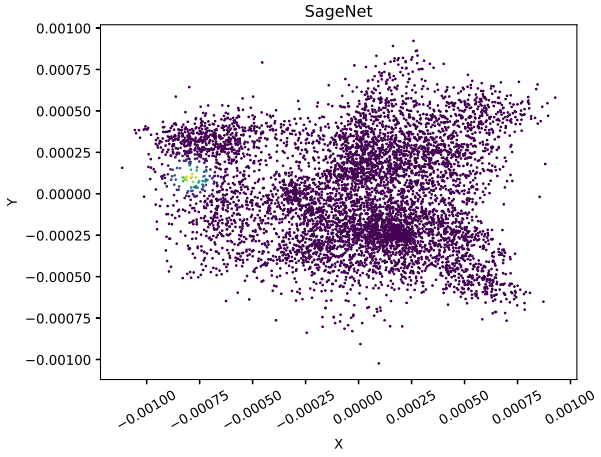

C

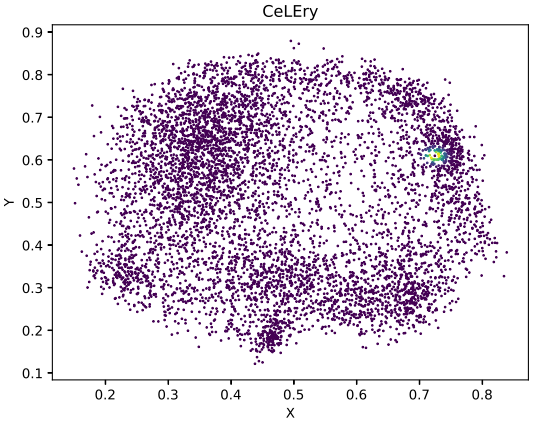

Supplementary Figure 4. Weight matrix of random cell’s neighbors for cell-type co-localization analysis using spatialDM. A, Weight matrix based on our method's reconstructed locations. B, Weight matrix based on SageNet’s reconstructed locations. C, Weight matrix based on by CeLEry’s reconstructed locations.

### 2. Supplementary Tables

Supplementary Table 1. Average spearman’s rank correlation coefficient on the mouse right brain cells derived by MERSCOPE.

| Method | Spearman's ρ |
| --- | --- |
| **Ours** | **0.766** |
| SageNet | 0.492 |
| CeLEry | 0.672 |
| novoSparc | 0.341 |
| Tangram | 0.137 |

Supplementary Table 2. Contingency table of detected cell-type co-localizations in reference datasets and SC sample that spatially reconstructed using our method.

| Detected co-localizations in reference datasets | Detected co-localizations in SC sample | |
| --- | --- | --- |
|  | Detected | Not detected |
| Reference support > 1 | 42 | 25 |
| Reference support ≤ 1 | 23 | 100 |

Supplementary Table 3. Contingency table of detected cell-type co-localizations in reference datasets and SC sample that spatially reconstructed using CeLEry.

| Detected co-localizations in reference datasets | Detected co-localizations in SC sample | |
| --- | --- | --- |
|  | Detected | Not detected |
| Reference support > 1 | 42 | 25 |
| Reference support ≤ 1 | 35 | 88 |

Supplementary Table 4. Contingency table of detected cell-type co-localizations in reference datasets and SC sample that spatially reconstructed using SageNet.

| Detected co-localizations in reference datasets | Detected co-localizations in SC sample | |
| --- | --- | --- |
|  | Detected | Not detected |
| Reference support > 1 | 38 | 29 |
| Reference support ≤ 1 | 26 | 97 |

Supplementary Table 5. Average spearman’s rank correlation coefficient for all benchmarking scenarios by setting the *m* as 21 (Eq. 3 in the Methods).

| Training ST dataset | Testing dataset | Spearman's ρ |
| --- | --- | --- |
| SeqFISH mouse embryo1 L1 | SeqFISH mouse embryo1 L2 | 0.89 |
|  | SeqFISH mouse embryo2 L2 | 0.634 |
|  | SeqFISH mouse embryo3 L2 | 0.532 |
| Stereo-Seq mouse embryo | SeqFISH mouse embryo1 L2 | 0.249 |
|  | SeqFISH mouse embryo2 L2 | 0.464 |
|  | SeqFISH mouse embryo3 L2 | 0.288 |
| 10X Visium human breast S06 | 10X Visium human breast S08 | 0.1 |
| MERSCOPE left mouse brain | MERSCOPE right mouse brain | 0.775 |
