## Supplementary material for "CellContrast: Reconstructing Spatial Relationships in Single-Cell RNA Sequencing Data via Deep Contrastive Learning": Description of Supplementary Data

### Description of Additional Supplementary Files

**Supplementary Dataset 1**: Benchmarking results of spatial reconstruction for mouse gastrulation cells (Embryo 1 L2) using single-cell ST reference.

**Supplementary Dataset 2**: Benchmarking results of spatial reconstruction for mouse gastrulation cells (Embryo 2 L2) using single-cell ST reference.

**Supplementary Dataset 3**: Benchmarking results of spatial reconstruction for mouse gastrulation cells (Embryo 3 L2) using single-cell ST reference.

**Supplementary Dataset 4**: Benchmarking results of spatial reconstruction for mouse gastrulation cells (Embryo 1 L2) using array-based ST reference.

**Supplementary Dataset 5**: Benchmarking results of spatial reconstruction for mouse gastrulation cells (Embryo 2 L2) using array-based ST reference.

**Supplementary Dataset 6**: Benchmarking results of spatial reconstruction for mouse gastrulation cells (Embryo 3 L2) using array-based ST reference.

**Supplementary Dataset 7**: Benchmarking results of human breast cells.

**Supplementary Dataset 8**: Benchmarking results of mouse brain cells.
